# Giant KASH proteins and ribosomes synergistically establish cytoplasmic biophysical properties *in vivo*

**DOI:** 10.1101/2025.01.10.632479

**Authors:** Xiangyi Ding, Hongyan Hao, Daniel Elnatan, Patrick Neo Alinaya, Shilpi Kalra, Abby Kaur, Sweta Kumari, Liam J. Holt, G. W. Gant Luxton, Daniel A. Starr

## Abstract

Understanding how cells control their biophysical properties during development remains a fundamental challenge. While cytoplasmic macromolecular crowding affects multiple cellular processes in single cells, its regulation in living animals remains poorly understood. Using genetically encoded multimeric nanoparticles for *in vivo* rheology, we discovered that *C. elegans* tissues maintain distinct cytoplasmic biophysical properties that differ from those observed across diverse systems, including bacteria, yeast species, and cultured mammalian cells. We identified two conserved mechanisms controlling cytoplasmic macromolecular diffusion: ribosome concentration, a known regulator of cytoplasmic crowding, works in concert with a previously unknown function for the giant KASH protein ANC-1 scaffolding the endoplasmic reticulum. These findings reveal mechanisms by which tissues establish and maintain distinct cytoplasmic biophysical properties, with implications for understanding cellular organization across species.

**One-Sentence Summary:** Living tissues maintain unique intracellular biophysical properties under the control of cytoplasmic constraints and crowding.

## Introduction

The cytoplasm is a complex biomolecular environment consisting of a protein-rich aqueous phase (the cytosol), dynamic networks of cytoskeletal filaments, and membrane-bound organelles. Macromolecules occupy up to 40% of cytoplasmic volume, creating a crowded environment that fundamentally alters diffusion, chemical reactions, and phase separations within cells (*1–8*). This macromolecular crowding (referred to as ‘crowding’ from here on) influences essential cellular processes including protein folding, metabolism, and signal transduction (*9*, *10*). Beyond crowding effects, molecular movement in the cytoplasm is further restricted by physical barriers and interactions. These include steric hinderance from the cytoskeletal network and organelles, as well as specific binding interactions with macromolecular complexes or molecular tethers. Together, crowding and these physical constraints define the biophysical properties of the cytoplasm.

Despite their importance, the biophysical properties of the cytoplasm in living multicellular organisms remain poorly understood. Traditional approaches for studying cytoplasmic biophysical properties rely on passive microrheology (*9*, *11*), where the motion of non-biological tracer particles is used to infer cytoplasmic dynamics, viscosity, elasticity, and structure. However, such studies have primarily focused on isolated cultured cells. The application of this technique to multicellular tissues has been limited by challenges in delivering probes into cells without disrupting cellular function, resulting in most studies being restricted to early embryos (*12*). Thus, there is a significant gap in our knowledge of the mechanisms that regulate cytoplasmic biophysical properties in complex, multicellular organisms.

Here, we investigate cytoplasmic biophysical properties in living multicellular animals using genetically encoded multimeric nanoparticles (GEMs) (*2*). These bright fluorescent tracers of defined shape and size serve as rheological tools across diverse organisms, including bacteria, cultured mammal cells, and yeast (*2*, *5*, *13*). We adapted GEMs for expression in the hypodermis and intestine of *Caenorhabditis elegans*, combining *in vivo* rheology with genetic manipulations to uncover mechanisms controlling cytoplasmic crowding and constraints.

### Exceptional cytoplasmic constraints at mesoscale in *C. elegans* tissues

To investigate cytoplasmic biophysical properties in living tissue, we engineered *C. elegans* strains with single-copy cDNA insertions encoding EGFP-tagged 40 nm in diameter GEMs expressed in the hypodermis or the intestine using the *semo*-*1* or *ges-1* promoters, respectively (Fig. 1A) (*14*). These cytoplasmic nanoparticles were readily visible in both tissues (Fig. 1B, Movie S1 and S2), and transgenic animals exhibited normal development, fertility, and viability (Fig. S1). We analyzed GEM motion in the anterior body region between the pharynx and oocytes of adult worms, capturing single-particle trajectories at 50 Hz using spinning disk confocal microscopy (Fig. 1B). From thousands of trajectories per tissue (minimum 10 frames), we calculated ensemble mean squared displacement (MSD) and effective diffusion coefficients (D_eff_) (Figs. 1C-D).

**Fig 1.**
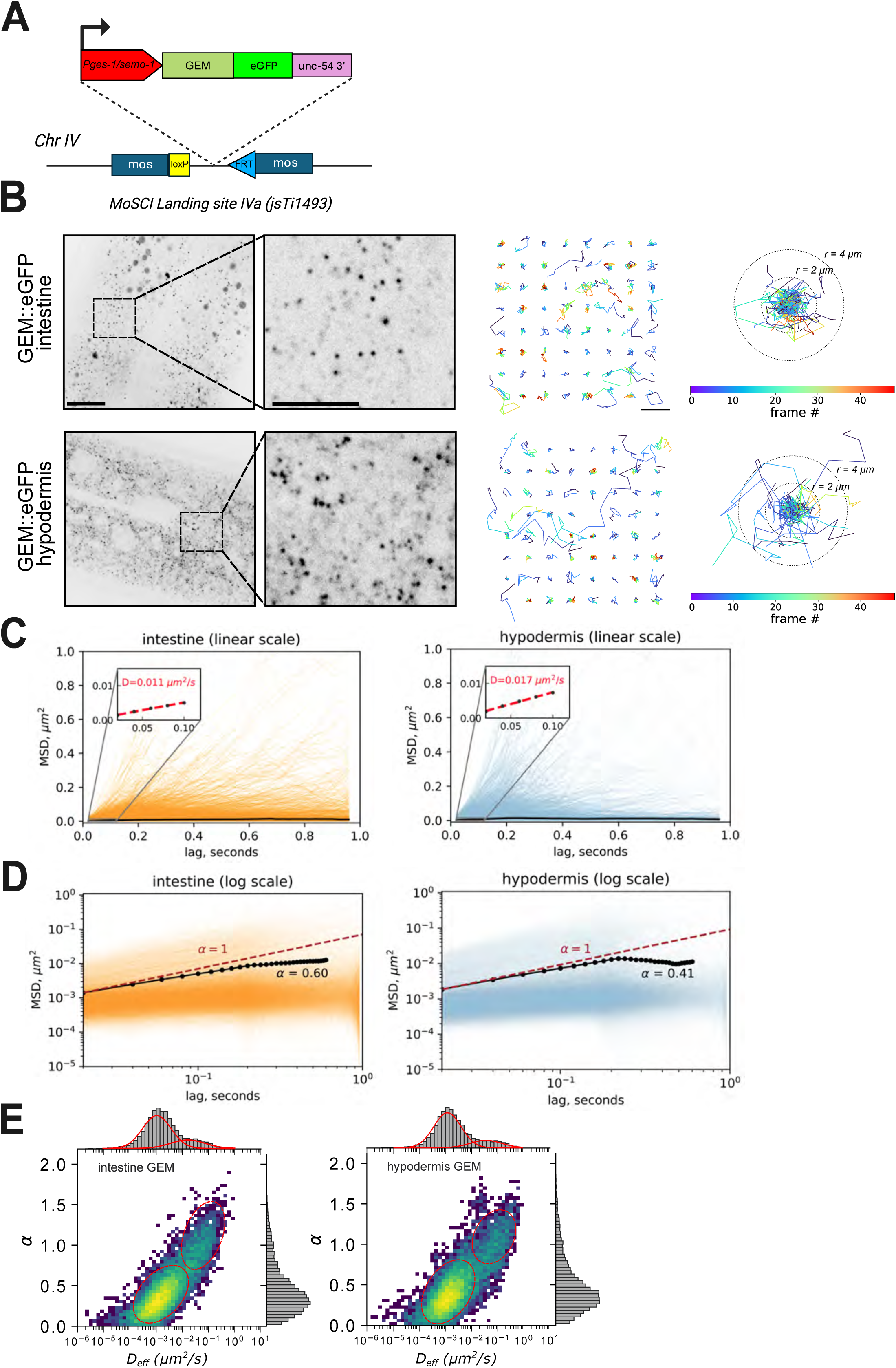
*C. elegans* tissues maintain highly crowded and constrained cytoplasmic environments. **A)** GEM expression cassette and genomic integration. A tissue-specific promoter (*Pges-1* for intestine or *Psemo-1* for hypodermis) drives the expression of a GEM-EGFP-encoding transgene integrated at the MosSCI landing site IVa (*jsTi1493*) on chromosome IV. **B)** Inverted grayscale spinning disc confocal images of GEMs in intestine (top) and hypodermis (bottom). Left: Representative fields of view (scale bar: 10 µm) with magnified insets (scale bar: 5 μm). Right panels display particle trajectories color-coded by time (scale bar: 4 μm, concentric circles indicate 2 μm and 4 μm radii). **C, D)** Mean squared displacement (MSD) analysis of GEM mobility in intestine and hypodermis shown in linear (C) with indicated diffusion coefficient D_eff_, and log-log (D) scale with indicated anomalous exponent (α). Individual particle trajectories (orange: intestine, blue: hypodermis) and ensemble average MSD (black lines) demonstrate restricted diffusion. Red dotted lines represent normal diffusion (α = 1). (E) 2D probability density histograms correlating D_eff_ with α for intestinal (left) and hypodermal (right) GEMs. Marginal distributions shown as histograms on axes. Red curves indicate two populations identified by Gaussian mixture modeling. Color intensity represents population density, revealing distinct mobility populations in both tissues.

GEMs in *C. elegans* tissues exhibited remarkably slow diffusion, with an ensemble D_eff_ below 0.02 µm^2^/s. This is more than 10-fold slower than what was previously observed across diverse single-cell systems, including budding and fission yeasts, bacteria, and cultured mammalian cells (HEK293, mouse embryonic fibroblasts) (*2*, *5*, *13*), suggesting that macromolecular diffusion in multicellular organisms is quite different than in single-cell models or mammalian tissue culture. The MSD curves obtained from both tissues showed slopes with an anomalous diffusion exponent (α) below 1 (indicated by the dashed line) (Fig. 1C-D), revealing sub-diffusive behavior and suggesting highly constrained GEM mobility. To understand the basis for this slow and constrained diffusion, we analyzed the relationship between D_eff_ and α. This revealed heterogeneous diffusion patterns consistent with cytoplasmic nonergodicity, with most particles displaying constrained movement. Using gaussian mixture modeling (*15*), we identified two distinct GEM populations: locally constrained particles and unconstrained but slow-moving particles (indicated by fitted curves, Fig. 1E). Notably, even the unconstrained GEMs diffused substantially slower than in single-cell systems(*2*, *16*). These findings reveal that *C. elegans* tissues maintain an exceptionally crowded and constrained cytoplasmic state.

### ANC-1 establishes size-dependent constraints on cytoplasmic diffusion

Giant Klarsicht/ANC-1/SYNE homology (KASH) proteins are massive molecular tethers that span the outer nuclear membrane and extend into the cytoplasm, where they interact with various cytoskeletal networks critical for cellular architecture (*17*, *18*). In *C. elegans*, the giant KASH protein ANC-1 prevents cytoplasmic ‘sloshing’ and maintains cellular integrity in the hypodermis (*19*). Similarly, disruption of ANC-1’s mammalian orthologs, nesprin-1 and −2, reduces cellular stiffness (*20*). We therefore hypothesized that ANC-1 constrains macromolecular motion, contributing to the remarkably slow diffusion we observed in *C. elegans* tissues. To test this hypothesis, we measured GEM dynamics in *anc-1(e1873)* mutants. GEM diffusion increased significantly in both intestinal and hypodermal tissues of *anc-1* mutants (Fig. 2A and S2, Movie S3 and S4). Analysis of individual trajectories revealed that loss of ANC-1 altered both the distribution and dynamics of GEM populations (Figs. 2C-F). The fraction of constrained particles decreased substantially, and bootstrap analyses showed significantly faster global diffusion compared to wild type (Figs. 2G-J).

**Fig 2.**
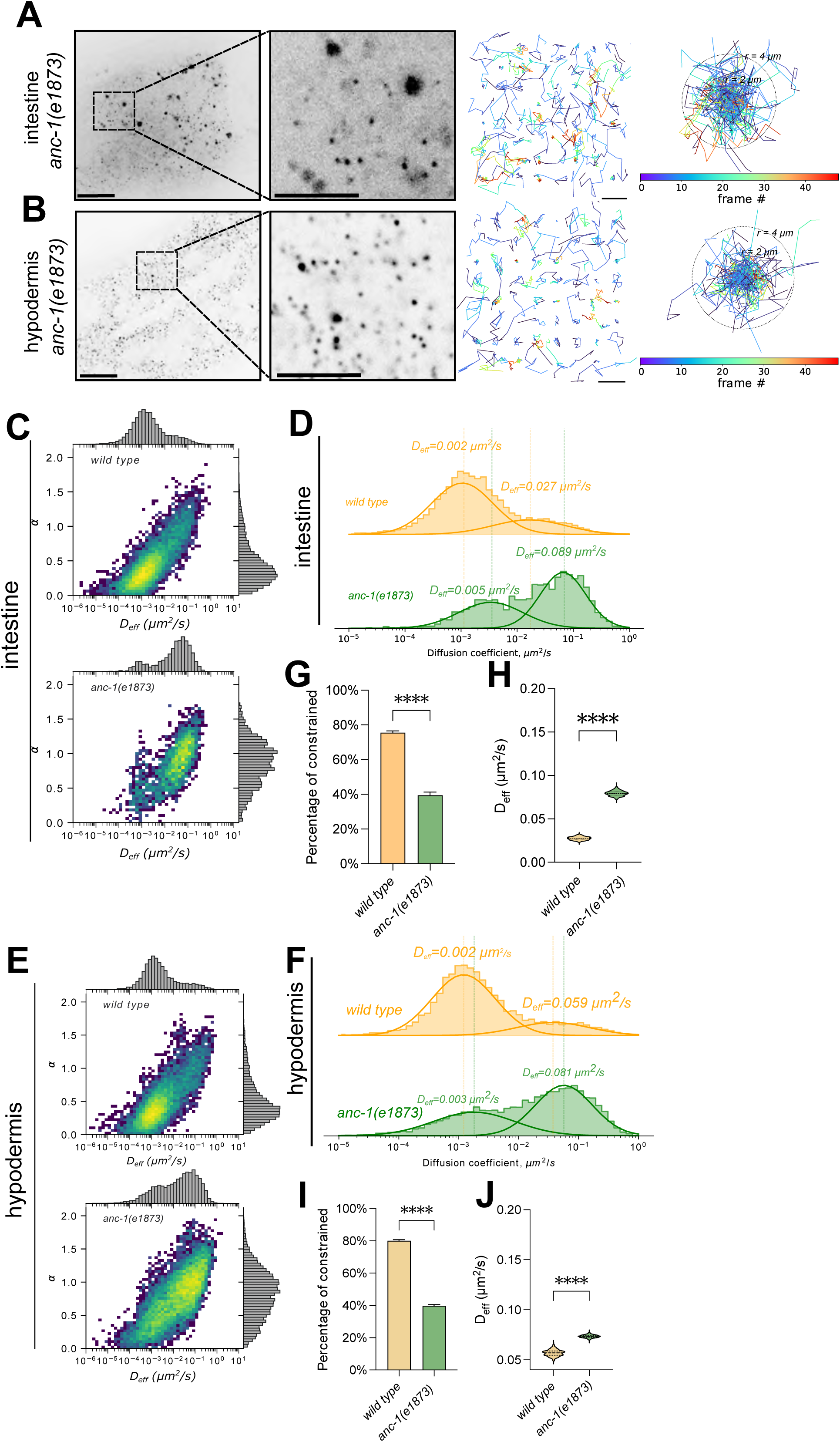
ANC-1 regulates cytoplasmic constraints in the hypodermis and intestine. **A, B)** Representative inverted grayscale spinning disc confocal images showing GEM mobility in intestinal (A) and hypodermal (B) cells in *wild type* or *anc-1(e1873)* animals. Left: Representative fields of view (scale bar: 10 µm) with magnified insets (scale bar: 5 μm). Right panels display particle trajectories color-coded by time (scale bar: 4 μm, concentric circles indicate 2 μm and 4 μm radii). **C, E)** Heat maps correlating D_eff_ with α for GEMs in intestinal (C) and hypodermal (E) cells in *wild type* or *anc-1(e1873)* animals. Marginal histograms show individual parameter distributions. Color intensity represents particle density. **D, F)** Gaussian mixture modeling (GMM) of GEM D_eff_ in intestine (D) and hypodermis (F) reveals distinct mobility populations. Orange: *wild type*; green: *anc-1(e1873)*. Dotted lines indicate population means. **G-J)** Bootstrap analysis of GMM parameters showing mean distributions for intestinal (G-H) and hypodermal (I-J) cells. **G, I)** Percentage of constrained GEMs. **H, J)** D_eff_ of unconstrained GEMs. Statistical significance assessed by Kruskal-Wallis test with Dunn’s multiple comparisons (****: p<0.0001).

To determine whether ANC-1’s effects were specific to large macromolecules, we performed fluorescence recovery after photobleaching (FRAP) experiments using cytoplasmic EGFP (∼4 nm in diameter) expressed in the hypodermis (*21*) (Fig. S2). Recovery times and diffusion rates were identical between wild type and *anc-1(e1873)* animals, demonstrating that ANC-1 specifically constrains the motion of larger macromolecules while allowing free diffusion of smaller proteins. These findings reveal an unexpected role for ANC-1 in establishing size-dependent constraints on cytoplasmic diffusion.

### Ribosomes and ANC-1 independently regulate mesoscale motion through crowding and constraint

Ribosomes are key determinants of cytoplasmic crowding in cultured yeast and mammalian cells (*2*, *22*, *23*). In addition, our forward genetic screen identified that knockdown of the ribosomal subunit *rps-15* caused a nuclear position defect in the syncytial hypodermis (Fig. S3). This led us to investigate how ribosome levels influence cytoplasmic GEM mobility *in vivo*. Using RNAi against two different ribosomal subunits (*rps-15* or *rps-18*), we found that ribosome depletion significantly increased GEM diffusion rates while maintaining the relative proportions of constrained and unconstrained populations (Figs. 3A-B, 3D, S3). The persistence of constrained GEMs in ribosome-depleted tissues suggests that ribosomes primarily affect cytoplasmic crowding rather than impose structural constraints.

**Fig. 3.**
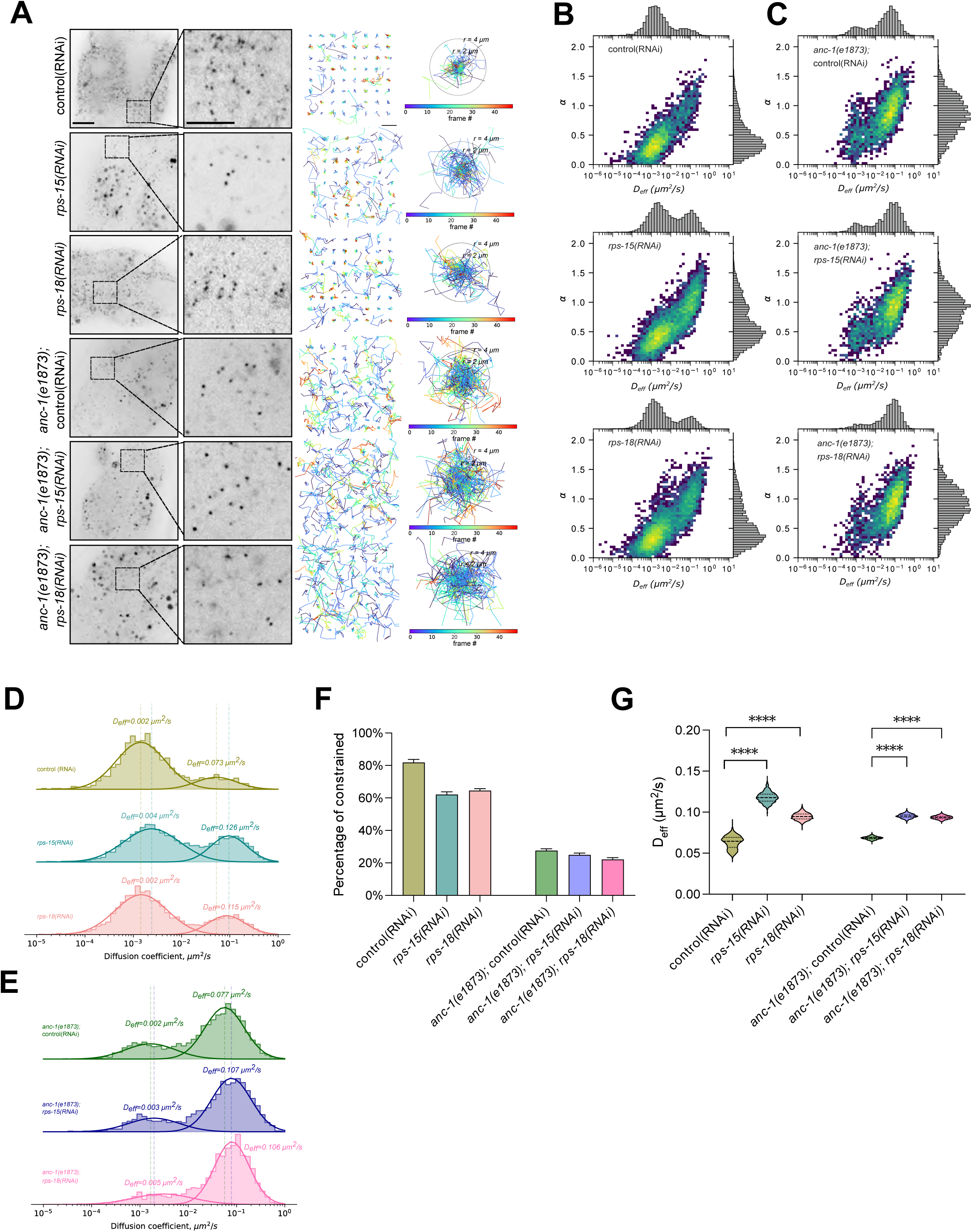
Ribosome depletion and ANC-1 loss affect cytoplasmic mobility through distinct mechanisms. **A)** Representative inverted grayscale spinning disc confocal images showing intestinal GEM mobility under control(RNAi), *rps-15(RNAi)*, and *rps-18(RNAi)* conditions. Left: Representative fields of view with magnified insets (scale bars: 10 μm and 5 µm, respectively). Right panels display particle trajectories color-coded by time (scale bar: 4 μm, concentric circles indicate 2 μm and 4 μm radii). **B)** Heat maps correlating D_eff_ with α for intestinal GEMs under ribosomal protein knockdown conditions. Marginal histograms show parameter distributions. **C)** Similar analysis in the *anc-1(e1873)* background with ribosomal protein knockdowns. **D)** GMM analysis of GEM *D_eff_* comparing control(RNAi) (olive), *rps-15(RNAi)* (blue), and *rps-18(RNAi)* (coral) knockdowns. Dotted lines indicate mean *D_eff_* values for control and *rps-15(RNAi)* populations. **E)** Similar analysis in the *anc-1(e1873)* background comparing control(RNAi) (green), *rps-15(RNAi)* (purple), and *rps-18(RNAi)* (pink) knockdowns. Dotted lines indicate mean *D_eff_* values for *anc-1(e1873);* control(RNAi) and *anc-1(e1873); rps-15(RNAi)* populations. **F-G)** Bootstrap analysis of GMM parameters comparing ribosomal protein knockdown conditions under *wild type* or *anc-1(e1873)* backgrounds. The percentage of constrained GEMs (F) with mean ± SD *D_eff_*values of unconstrained GEMs (G) are shown. Statistical significance was assessed by Kruskal-Wallis test with Dunn’s multiple comparisons (****: p<0.0001).

To investigate potential interactions between ribosome-mediated crowding and ANC-1-dependent constraint, we depleted ribosomes in *anc-1(e1873)* tissues (Fig.3A). Combined loss of ANC-1 and ribosomes produced striking effects: while the shift toward unconstrained GEMs was only marginally greater than in *anc-1* mutants alone, the diffusion rates of unconstrained GEMs dramatically increased compared to either single perturbation (Figs. 3C, 3E-G, and S3). These distinct phenotypic signatures reveal that ANC-1 and ribosomes control cytoplasmic macromolecular mobility through complementary mechanisms – ANC-1 by imposing structural constraints and ribosomes through crowding.

### The transmembrane α-helix of ANC-1 is essential for cytoplasmic constraints

ANC-1 is a large protein (∼800 kDa) composed of multiple conserved domains: N-terminal tandem calponin homology (CH) domains that bind actin, a central region containing six tandem repeats (RPs) of approximately 900 residues each predicted to form spectrin-like structures, and a C-terminal KASH domain with its transmembrane α-helix that localizes to the nuclear envelope and endoplasmic reticulum (ER) membranes (Fig. 4A) (*19*, *24*). Having established ANC-1’s role in regulating cytoplasmic constraints, we investigated its mechanism of action. Our findings revealed that ANC-1 maintains cytoplasmic integrity through a pathway independent of its canonical role in the linker of nucleoskeleton and cytoskeleton (LINC) complex. Previous work showed that the transmembrane α-helix and 6RPs are essential for organelle anchorage and cytoplasmic integrity, while the C-terminal KASH peptide and N-terminal actin-binding CH domains are dispensable(*19*). To precisely identify the ANC-1 domains critical for maintaining cytoplasmic constraints, we analyzed cytoplasmic GEM diffusion in mutants lacking specific regions: the KASH peptide (ΔKASH), the transmembrane α-helix plus KASH peptide (ΔTK), the 6RPs (Δ6RPs), or the N-terminal actin-binding CH domains (ΔCH) (Fig. 4A, S4). Our analysis revealed tissue-specific requirements for ANC-1 domains. In both tissues examined, *anc-1(ΔTK)* mutants showed disrupted cytoplasmic constraints comparable to *anc-1(e1873)* animals (Figs. 4B-C). In contrast, minimal effects on GEM diffusion were observed in *anc-1(ΔKASH), anc-1(Δ6RPs),* or *anc-1(ΔCH)* mutants (Fig. S4). In intestinal cells, GEMs diffused significantly faster in *anc-1(ΔTK)* mutants compared to mutants lacking other ANC-1 domains (Figs. 4D, 4F, 4H). However, in hypodermal cells, both *anc-1(ΔTK)* and *anc-1(ΔKASH)* mutants displayed enhanced particle mobility (Figs. 4E, 4G, 4I). This suggests that there are interesting tissue-specific differences in the role of ANC-1 modulating biophysical properties of the cytoplasm. These findings also demonstrate that the transmembrane α-helix of ANC-1 is essential for maintaining cytoplasmic constraints. Intriguingly, fewer spectrin-like repeats, which create extensible head-to-tail rod-like structures, are required for constraining macromolecules compared to anchoring organelles.

**Fig. 4.**
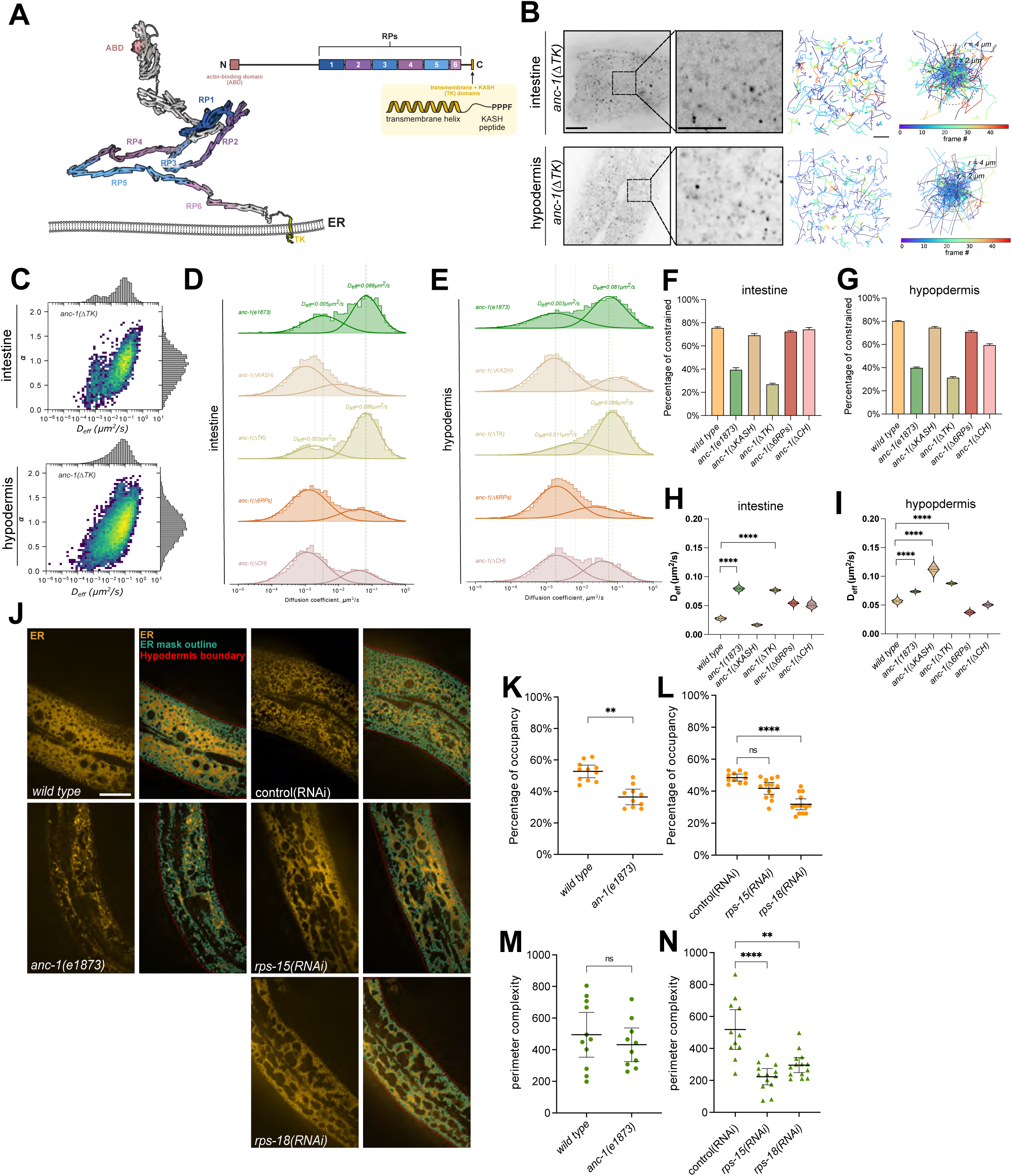
ANC-1’s transmembrane α-helix regulates cytoplasmic constraints and ER organization. **A)** AlphaFold structural prediction of ANC-1 showing the domain architecture, including the N-terminal actin-binding domain (ABD, pink), six repeat (RP) domains (RP1-RP6, alternating blue and purple), and C-terminal transmembrane (T) domain that spans both the ER and nuclear envelope membranes with the KASH (K) peptide extending into the perinuclear space. The linear domain diagram above shows the corresponding domain organization. **B)** Representative inverted grayscale spinning disc confocal images showing GEM mobility in intestinal and hypodermal tissues of *anc-1(ΔTK)* mutants. Left: Representative fields of view with magnified insets (scale bars: 10 μm and 5 µm, respectively). Right panels display particle trajectories color-coded by time (scale bar: 4 μm, concentric circles indicate 2 μm and 4 μm radii). **C)** Heat maps correlating D_eff_ and α for intestinal and hypodermal GEMs in *anc-1(ΔTK)* mutants. Marginal histograms show parameter distributions. **D-E)** GMM of GEM D_eff_ values comparing *anc-1(e1873)* (green) with *anc-1(ΔTK)* mutants (beige) in intestine (D) or hypodermis (E). Dotted lines indicate population means. **F-I)** Bootstrap analysis of GMM parameters across genotypes. Percentage of constrained GEMs in intestine (F) and hypodermis (G). D_eff_ values of unconstrained GEMs in intestine (H) and hypodermis (I). **J)** Representative spinning disc confocal images of hypodermal ER organization. Left: ER signal; Right: Thresholded images showing ER outline (cyan) and hypodermal boundary (red). Scale bar: 20 μm). **K-N)** ER network analysis: ER occupancy percentage (K, L) and perimeter complexity (M, N) across genotypes. Statistical significance was assessed by Kruskal-Wallis test with Dunn’s multiple comparisons (*: p≤0.05, **: p≤0.01, ***: p≤0.001, ****: p≤0.0001, ns: p>0.05)

### ANC-1 and ribosomes maintain ER architecture through distinct structural and crowding mechanisms

The ANC-1 transmembrane α-helix is essential both for ER localization (*19*) and maintenance of cytoplasmic constraints, suggesting ANC-1 might function by maintaining ER structure and organization. Given that ribosomes are major determinants of crowding and associate extensively with the ER, we investigated how both ANC-1-dependent cytoplasmic constraint and ribosome-dependent cytoplasmic crowding contribute to ER organization. *anc-1(e1873)* and *rps-18(RNAi)* mutants exhibited distinct defects in ER organization. Ribosome depletion resulted in large, interconnected cytoplasmic vacancies in the ER network, while *anc-1(e1873)* animals showed increased ER fragmentation compared to *rps-18(RNAi)* animals (Fig. 4J). Quantitative analysis revealed decreased ER occupancy (See methods) in both conditions (Fig. 4K, 4L), though network complexity (See methods) was reduced only in *rps-18(RNAi)* animals (Fig. 4N). Despite the ER morphology defects observed in *rps-18(RNAi)* animals, cytoplasmic integrity remained stable during locomotion, contrasting with the pronounced ER ‘sloshing’ defect observed in *anc-1(e1873)* mutants (Movie S5) (*19*). Additionally, ribosome levels, measured by RPS-18::GFP intensity, remained unchanged in *anc-1(e1873)* mutants (Fig. S5).

These findings support a model where ANC-1 and ribosomes regulate cytoplasmic biophysical properties through distinct but complementary mechanisms. Ribosomes function as crowding agents, like their role in other systems (*2*, *5*), and through this crowding maintain ER morphology. In contrast, ANC-1 provides structural support through its spectrin-like repeats, which are targeted to the ER via its transmembrane α-helix. This scaffolding role, characteristic of spectrin proteins (*19*, *25*), may allow the ANC-1 supported ER to constrain macromolecular complexes and maintain cytoplasmic integrity. Together, this dual mechanism ensures both the structural stability and proper organization of the ER network while maintaining appropriate cytoplasmic biophysical properties *in vivo*.

### Summary

Our study reveals fundamental mechanisms controlling cytoplasmic biophysical properties in living animal tissues. Using GEMs, we discovered that *C. elegans* tissues maintain an exceptionally constrained cytoplasmic state compared to other biological systems. This tight control over cytoplasmic properties emerges from two distinct but complementary mechanisms: ribosome-mediated crowding and ANC-1-dependent structural constraints. We show that the giant KASH protein ANC-1 plays a previously unrecognized role in establishing size-dependent constraints on cytoplasmic diffusion through its association with the ER network. This function requires ANC-1’s transmembrane α-helix but is independent of its canonical role in nuclear anchoring. Simultaneously, ribosomes act as critical crowding agents that help maintain ER network organization, revealing how cells use both structural and crowding mechanisms to control their internal architecture. These findings establish a new paradigm for understanding cellular organization, where cells employ both passive (crowding) and active (structural) mechanisms to maintain proper cytoplasmic properties. This dual control system may represent a conserved strategy allowing tissues to establish and maintain distinct biophysical environments appropriate for their specialized functions. Future studies can build on this framework to investigate how different cell types tune their cytoplasmic properties during development and in response to environmental challenges.

## Acknowledgments

We thank members of the Holt and Starr-Luxton labs for helpful discussions, Dr. Thomas Wilkop and the MCB Light Imaging Facility, Wormbase, and Dr. Min Han (University of Colorado, Boulder) in whose lab the forward genetic screen was performed.

## Author contributions

Conceptualization: XD, HH, LJH, GWGL, DAS

Methodology: XD, HH, DE, LJH, GWGL, DAS

Investigation: XD, HH, DE, PA, SK, AK, SK

Visualization: XD, DE

Funding acquisition: LJH, GWGL, DAS

Project administration: GWGL, DAS

Supervision: GWGL, DAS

Writing – original draft: XD, GWGL, DAS

Writing – review & editing: XD, HH, DE, LJH, GWGL, DAS

## Data and materials availability

All data are available in the main text or the supplementary materials.

## Supplementary Materials

### Materials and Methods

#### Chemicals and Molecules

QIAprep Spin Miniprep Kit for DNA purification was obtained from Qiagen (Hilden, Germany). Restriction enzyme SapI and T4 DNA ligase were obtained from New England Biolabs (Ipswich, MA). The GEM gene block was synthesized by Integrated DNA Technologies (Coralville, IA). RNAi feeding vectors and bacterial strains were obtained from the Ahringer RNAi library (Source Bioscience, Nottingham, UK).

#### GEM Plasmid Construction and Strain Generation

The 40nm GEM expression constructs were generated using the encapsulin protein sequence from *Pyrococcus furiosus* (PDB ID: 2E0Z). The sequence consisted of the encapsulin open reading frame fused to EGFP, the *unc-54* 3’ UTR, and flanking *Sap*I restriction sites. The sequence was codon-optimized for *C. elegans* expression and included artificial introns to enhance its expression (Fig. S2). The optimized sequence was commercially synthesized as a gene block (Integrated DNA Technologies, Coralville, IA),

Two tissue-specific promoters were used for expression: the *semo-1* promoter (2.9 kb) for hypodermal expression and the *ges-1* promoter (2 kb) for intestinal expression. Primers were designed with complementary overhands to both the GEM gene block and the backbone vector pLF3FShC(*26*). The final constructs were assembled using Golden Gate cloning with the *Sap*I restriction enzyme and T4 DNA ligase, generating plasmids pSL848 (*p_ges-1_*) and pSL850 (*p_semo-1_*). Plasmid sequences were verified by Sanger sequencing.

*C. elegans* strains were maintained on nematode growth media (NGM) plates seeded with *E. coli* strain OP50(*27*, *28*). Some strains were obtained from the Caenorhabditis Genetics Center, funded by the National Institutes of Health Office of Research Infrastructure Programs (P40 OD010440). Single-copy transgenic strains were generated using FLP recombinase-mediated cassette exchange (RMCE)(*26*). Plasmids pSL848 and pSL850 (50 ng/μL) were microinjected into the gonads of the RMCE-compatible strain NM5179 to generate strains UD803 and UD838. All strains used in this study are listed in Supplemental Table S1.

The endogenous GFP11 tag was introduced into the *rps-18* locus using CRISPR/Cas9 genome editing (*29*). A crRNA (caagatgtcgttgatcattc) was used to guide Cas9 to the target site. For homology-directed repair, a DNA repair template containing GFP11 flanked by homology arms was used (IDT HDR ssDNA). The repair template sequence was: CTAAATTTTTTATTTTTCAGGGTCATAAAATCACGACAAGATGAGAGATCACATGGTTCTTCATGAATATGTAAATGCAGCTGGAATTACAGAACTCGGCTCAGGATCTGGTTCTTCAGCTGGTTCGTTGATCATCCCAGAGAAATTCCAGCACATTCATCGTGTGATGAACACCAACATCGAT. Successful genome editing was confirmed by PCR and sequencing.

#### RNAi-based genetics

Expression levels of the intestinal GEM strain (UD803) were suitable for single-particle tracking, but the hypodermal strain (UD838) had expression levels too high for efficient particle tracking. To achieve optimal expression levels in the hypodermal strain, animals were fed bacteria expressing dsRNA targeting *gfp* to reduce GEM-EGFP fusion protein levels(*30*, *31*).

RNAi experiments were conducted by feeding HT115(DE3) *E. coli* transformed with L4440-based vectors expressing double-stranded RNA. RNAi clones were obtained from the Ahringer RNAi library (Source Bioscience, Nottingham, UK)(*30*), and verified by Sanger sequencing before use. Bacteria were seeded onto NGM plates according to established protocols(*30*). Synchronized L2-L3 stage worms were transferred to RNAi feeding plates and maintained at 23° C for 48 hours until reaching the young adult stage prior to imaging.

A forward genetic screen for nuclear position defects similar to *anc-1* mutants was performed using RNAi feeding. Single adult *kuIs54[sur-5::gfp]* (MH1870; gift of Min Han, University of Colorado, Boulder) animals with GFP-marked nuclei(*32*) were placed on plates containing *E. coli* expressing RNAi clones from chromosome I of the Ahringer library (Source Bioscience)(*30*). Offspring were examined for nuclear positioning defects in the syncytial hypodermis using a Leica MLZIII fluorescent dissecting stereo microscope with a 2x objective (Leica, Deerfield, IL). Screening of all chromosomes. The clone yielded two hits: multiple clones targeting *anc-1 and* clone F36A2.6 against *rps-15*. The nuclear anchorage phenotypes of *rps-15(RNAi)* and *rps-18(RNAi)* animals were quantified using previously described methods(*33*).

#### Spinning Disc Confocal Microscopy

Imaging was performed on a Nikon Ti2 microscope (Nikon Instruments, Melville, NY) equipped with a Yokogawa CSU-X1 confocal scanner unite (Yokogawa Electric Corporation, Tokyo, Japan), and a Hamamatsu ORCA-Flash4.0 LT3 Digital sCMOS camera (Hamamatsu Photonics, Shizuoka, Japan), using a Nikon Plan Apo l 100X oil immersion objective (N.A. 1.45). Images were acquired at a resolution of 1060 x 1568 pixels at 0.065 µm/pixel. For visualization of GEMs, samples were illuminated using a Nikon 488 nm laser at 100% power and imaged with an exposure time of 20 ms per frame under continuous acquisition mode (i.e., no inter-frame interval). The ER was visualized using Nikon 561 nm laser at 50% power with a 100 ms exposure time. Ribosome was visualized using Nikon 488 nm laser at 30% power with 50 ms exposure time. Image acquisition was performed using Nikon Elements software. Images for GEMs and ER are processed and analyzed in napari GUI with GEM Detection and Motion Analysis or ER Occupancy and Complexity Analysis (See methods). The ribosome marker intensity and colocalization with ER marker were measured and analyzed in FIJI/ImageJ manually.

#### Cytoplasmic EGFP FRAP Experiments and Analysis

*C. elegans* L4 larvae expressing cytoplasmic EGFP under the *sur-5* promoter in wild type or *anc-1(e1873)* backgrounds (strains UD1053 and UD1058, respectively) were grown at 25° C for 24 hours before imaging. FRAP experiments were performed on a Zeiss 980 Laser Scanning Microscope at the University of California, Davis, MCB Light Microscopy Imaging Facility equipped with Airyscan 2 detection system (Carl Zeiss AG, Jena, Germany). Young adults were mounted on 2% agarose pads containing 7 μL of 1 mM tetramisole hydrochloride 5 minutes before imaging. The hypodermis between the pharynx and germline was visualized using a Zeiss 63X oil immersion objective (N.A. 1.40) with an additional 8X zoom. Images were acquired at 512 × 512 pixels (bidirectional pixel dwell time: 0.34 μs, pinhole: 1 Airy unit). Two 3.71 μm × 3.85 μm regions of interest (ROIs) were defined for photobleaching (Zeiss 488 nm laser at 100% power with a Gaussian profile, 20 iterations, 0.5 ms/iteration), post-bleach recovery monitoring (12 seconds at 103.22 ms intervals). Photobleaching achieved 60-70% reduction in fluorescence intensity.

FRAP analysis followed modified protocols from(*34*). Briefly, fluorescence intensities were corrected for photobleaching, baseline-adjusted to initial post-bleach intensity, and normalized to pre-bleach fluorescence. Recovery curves were fitted to a single exponential equation I(t) = I_max_(1-e(-kt)) using PRISM 9 (Dotmatics, Boston, MA), where I(t) is fluorescence intensity at time t, I_max_ is maximal recovery intensity, and k is the recovery rate constant. Half-life (t_1/2_) was calculated for each recovery curve.

#### GEM Detection and Motion Analysis

GEM particle detection was performed by modeling diffraction-limited fluorescent spots as two-dimensional (2D) Gaussians above a spatially varying local background(*35*). Initial spot detection used an 11 x 11 pixel ROI, fitted to a 2D Gaussian function using maximum-likelihood estimation for Poisson-distributed data (*36*). The spot detection and localization algorithms were implemented in a custom Python/C program (spotfitlm) (*37*). Particle tracking was performed using TrackPy v0.6.4 (*38*) with the ‘link’ function (search range=4.2). Trajectories shorter than 10 frames (0.2 seconds) were excluded from MSD analysis. For each trajectory, the *D_eff_* was calculated by linear fitting of the first 5 points of the MSD vs. time plot using ‘numpy.polyfit’. The anomalous coefficient α was determined by fitting a line through the log(MSD) vs. log(time) plots, using ‘scipy.optimize.least_squares’ with the “soft-L1” method minimizing the weighted residual function:

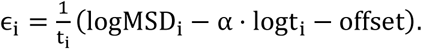

Diffusion coefficient distributions were analyzed after log-transformation, which yielded normal distributions. Two-component Gaussian mixture models were fitted using ‘scikit-learn.mixture.GaussianMixturè to resolve sub-populations. Mean (μ) and variance (σ^2^) parameters were converted from their log-transformed values (denoted by subscript) using standard log-normal distribution formulae

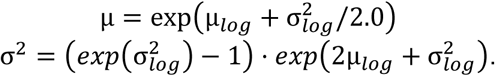

Parameter distributions were estimated by bootstrapping, resampling each dataset with replacement 1,000 times for Gaussian mixture model analysis.

#### ER Occupancy and Complexity Analysis

ER fluorescence images were initially processed using background subtraction with an iterative wavelet transform to remove low-frequency background(*39*). The original discrete wavelet transform was replaced by an undecimated wavelet transform (à trous) algorithm using a cubic B-spline kernel. We computed five levels of detail coefficients and ran the algorithm for a maximum of 25 iterations.

The background-subtracted images were then denoised and smoothed by minimizing a convex objective function:

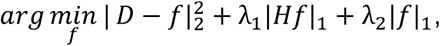

Where the first term enforced data fidelity, the second term (scaled by *λ*_1_) controlled smoothness and continuity, and the third term (scaled by *λ*_2_) controlled output sparsity. *D* represented the background-subtracted image, with *f* as the solution variable. We used the alternating direction method of multiplier algorithm(*40*) to efficiently obtain solutions to the sparsity-inducing L1-norm (denoted by | ⋅ |_1_). *H* represented a second-order finite difference operator. Computation was GPU-accelerated using PyTorch(*41*).

We implemented the background-subtraction and denoising workflow as a custom Napari widget(*42*, *43*). Denoising parameters were set to ρ = 0.1, λ_1_ = 10 − 40, λ_2_ = 10 − 40. Denoised images were converted to binary ER masks using Otsu’s method in scikit-image (*44*). Worm body masks were manually drawn in Napari. For 2D ER morphology quantification, we calculated ER occupancy defined by the ratio ER mask / worm body mask, and we calculated ER complexity defined by the ratio *P*^2^ / (4πA)(*45*), where *P* and *A* represented the shape’s perimeter and area, respectively.

**Fig. S1.**
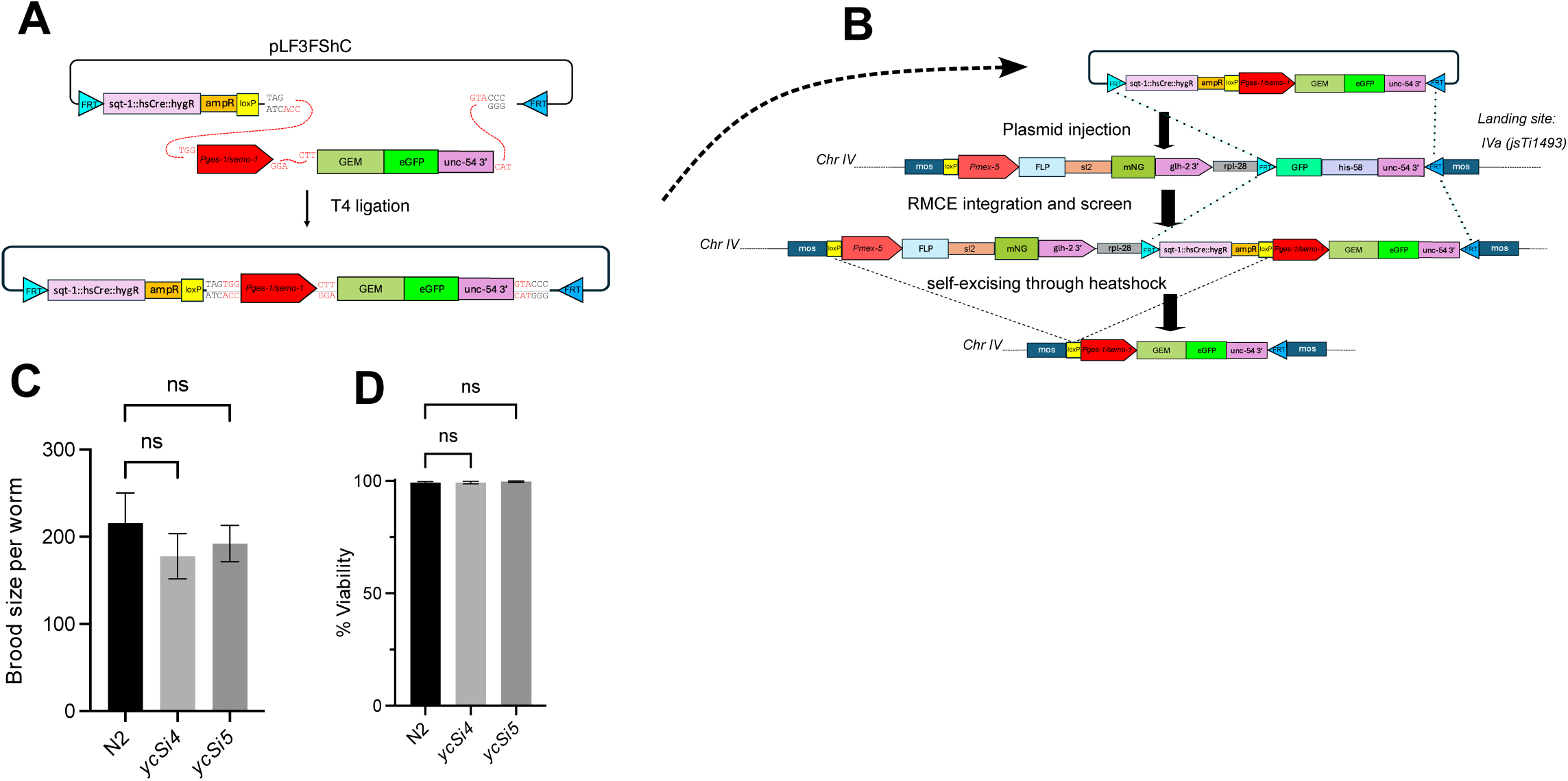
GEM transgene integration maintains normal *C. elegans* fertility and viability. **A)** Schematic showing GEM expression cassette assembly in the pLF3FShC plasmid using T4 ligation. **B)** Single-copy transgene integration workflow: GEM cassette injected into worms containing the MosSCI landing site IVa (jsTi1493) followed by RMCE-mediated integration and heat shock-induced cassette removal (see Materials and Methods and Nonet, 2020 (*26*)). **C-D)** Phenotypic analysis comparing wild type with N2 allele and GEM expressing strains with single copy GEM insertion allele *ycSi4* (intestinal) or *ycSi5* (hypodermal): Average brood size (C) and embryonic viability (D). Data shown as mean ± 95% confidence interval (CI). Statistical significance was assessed by a One-way ANOVA with Dunnett’s test (ns: p>0.05).

**Fig. S2.**
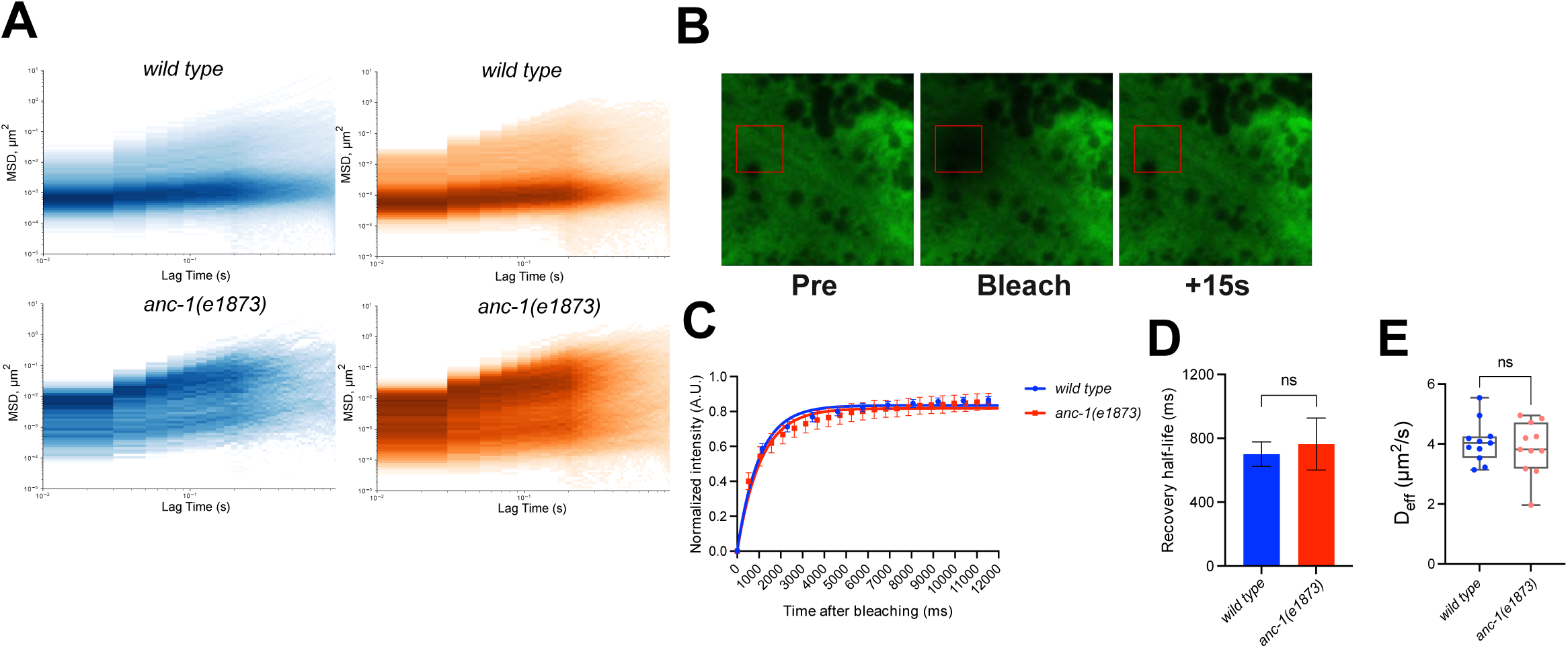
ANC-1 specifically affects mesoscale but not nanoscale cytoplasmic diffusion. **A)** Heat maps showing MSD vs. lag time for intestinal (blue) and hypodermal (orange) GEMs in *wild type* and *anc-1(e1873)*. Color intensity indicates trajectory density. **B-E)** FRAP analysis of cytoplasmic EGFP in hypodermal cells: (B) Representative time-lapse series showing pre-bleach, bleach, and 15-second post-bleach recovery (bleached region indicated by red box). (C) Fluorescence recovery curves comparing wild type and *anc-1(e1873)*. Points show normalized intensity; lines show exponential fits. (D) Recovery half-times and (E) calculated *D_ef_*_f_ values. Data presented as mean ± 95% CI. Statistical significance was assessed by Kolmogorov-Smirnov test (ns: p > 0.05).

**Fig. S3.**
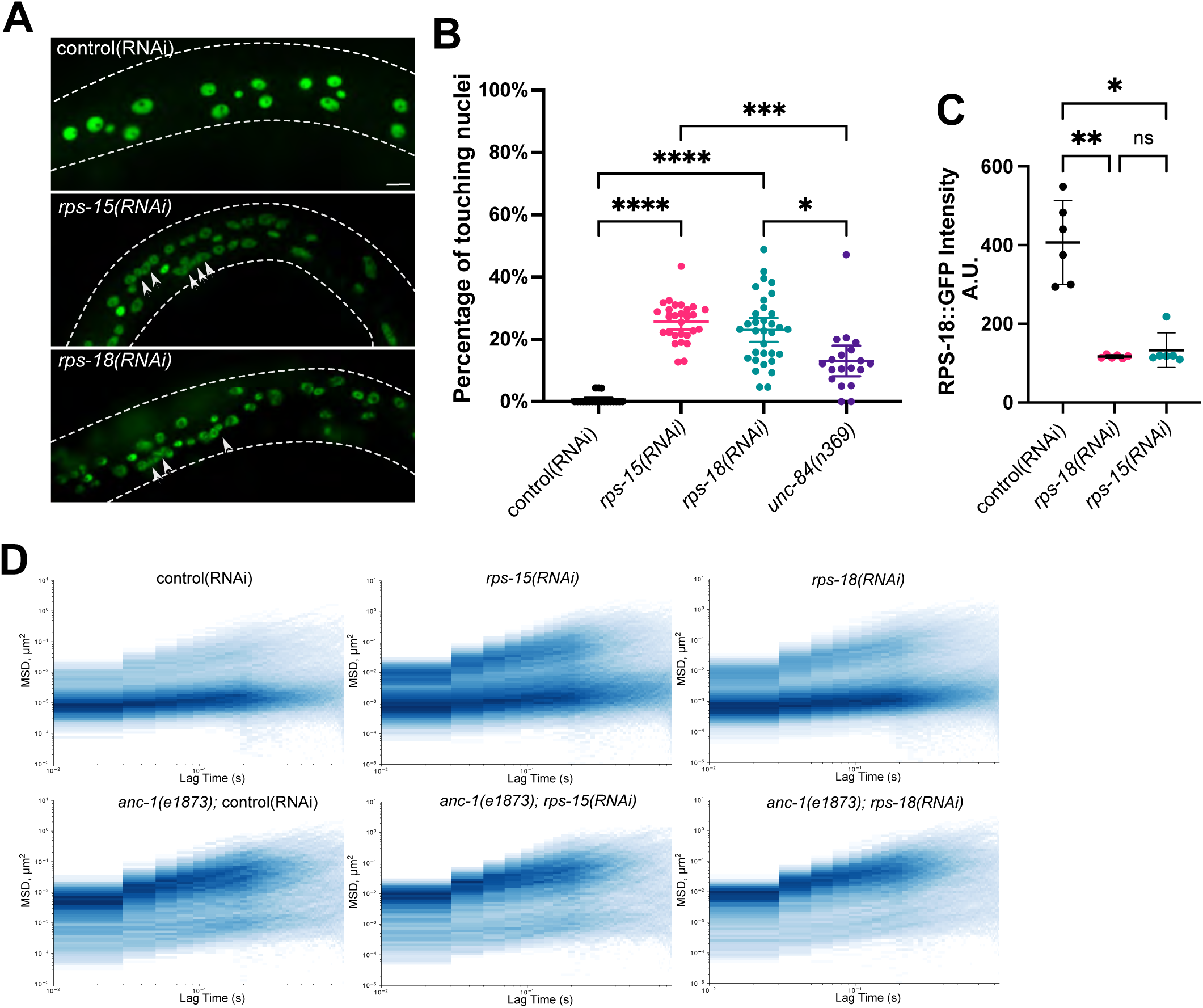
RPS-15 and RPS-18 affect both cytoplasmic crowding and hypodermal nuclear positioning. **A)** Representative spinning disc confocal images of EGFP-labeled hypodermal nuclei after RNAi treatment. Arrowheads: defective nuclear anchoring; dotted lines: worm outline. Scale bar: 10 μm. **B)** Nuclear positioning defects quantified as percentage of touching nuclei per worm under different RNAi conditions. **C)** RPS-18::GFP fluorescence intensity measurements showing protein depletion after *rps-15(RNAi).* Data presented as mean ± 95% CI. **D)** Heatmaps of MSD vs. lag time for GEMs in *wild type* and *anc-1(e1873)* backgrounds under different RNAi treatments. Statistical significance was assessed by Kruskal-Wallis test with Dunn’s comparisons (*: P<0.05, **: P<0.01, ***: P<0.001, ****: P<0.0001, ns: p>0.05).

**Fig. S4.**
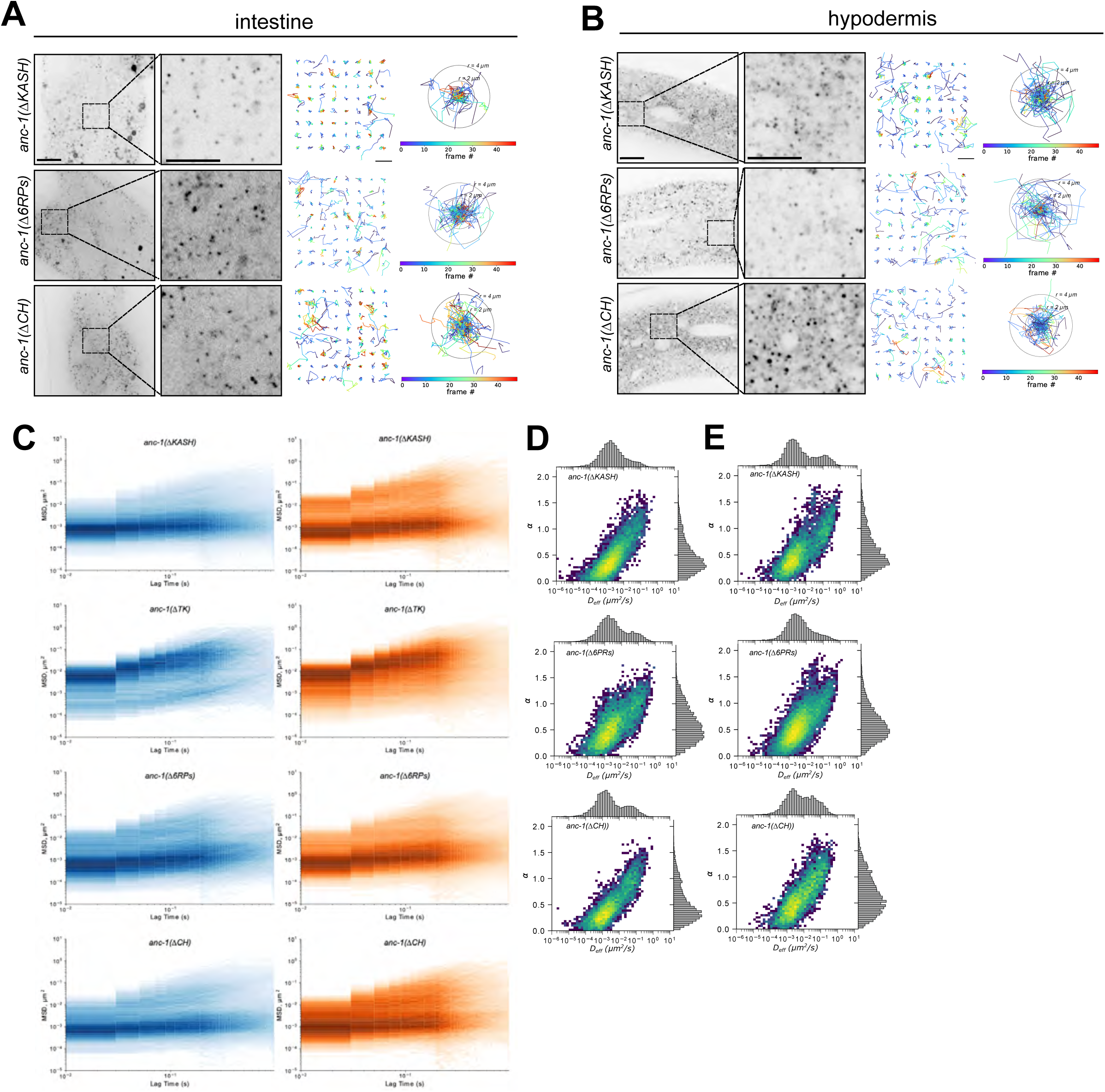
ANC-1’s transmembrane α-helix is essential for cytoplasmic constraint regulation. **A-B)** Representative inverted grayscale spinning disc confocal images showing GEM mobility in intestinal (A) and hypodermal (B) tissues of ANC-1 domain deletion mutants (ΔKASH, Δ6RPs, and ΔCH). Left: Representative fields of view with magnified insets (scale bars: 10 μm and 5 µm, respectively). Right panels display particle trajectories color-coded by time (scale bar: 4 μm, concentric circles indicate 2 μm and 4 μm radii). **C)** Heat maps of MSD vs. lag time in intestinal (blue) and hypodermal (orange) tissues for each domain deletion. **D-E)** Heat maps correlating *D_eff_* values with α for intestinal (D) and hypodermal (E) GEMs across mutants. Marginal histograms are shown.

**Fig. S5.**
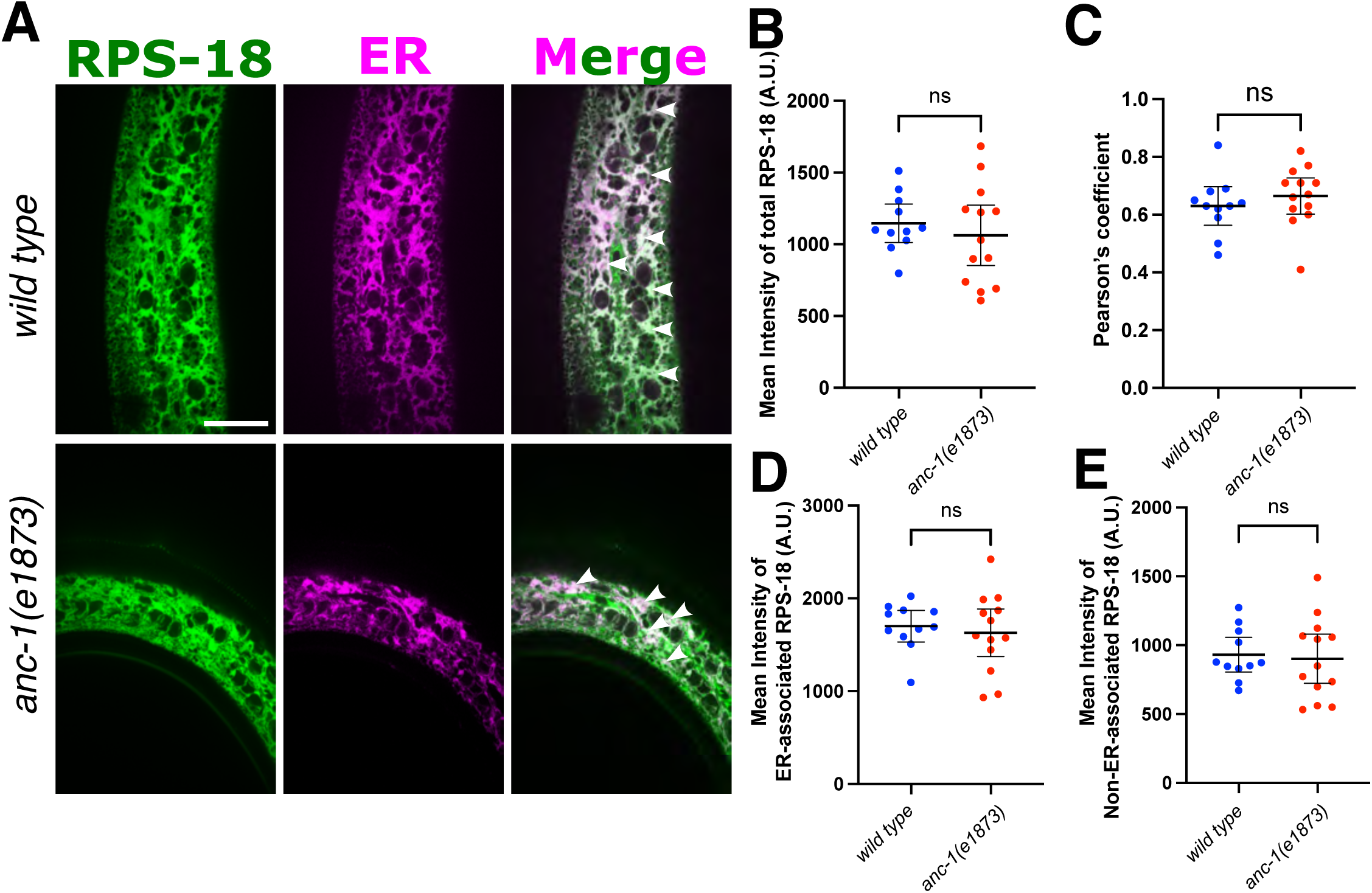
ANC-1 loss does not affect ribosome levels or ER association. **A)** Representative spinning disc confocal images of hypodermal tissue showing RPS-18 (green), ER (magenta), and merged signals in wild type and *anc-1(e1873)* animals (scale bar: 20 μm). Arrowheads emphasize the colocalized ER and ribosome. **B)** Total hypodermal RPS-18 intensity measurements. **C)** RPS-18/ER colocalization measured by Pearson’s correlation. **D-E)** Mean intensity of ER-associated (D) and non-ER-associated (E) RPS-18. Intensity is represented by arbitrary unit (A.U.) Data presented as mean ± 95% CI. Statistical significance was assessed by Kolmogorov-Smirnov test (ns: p>0.05).

**Fig. S6.**
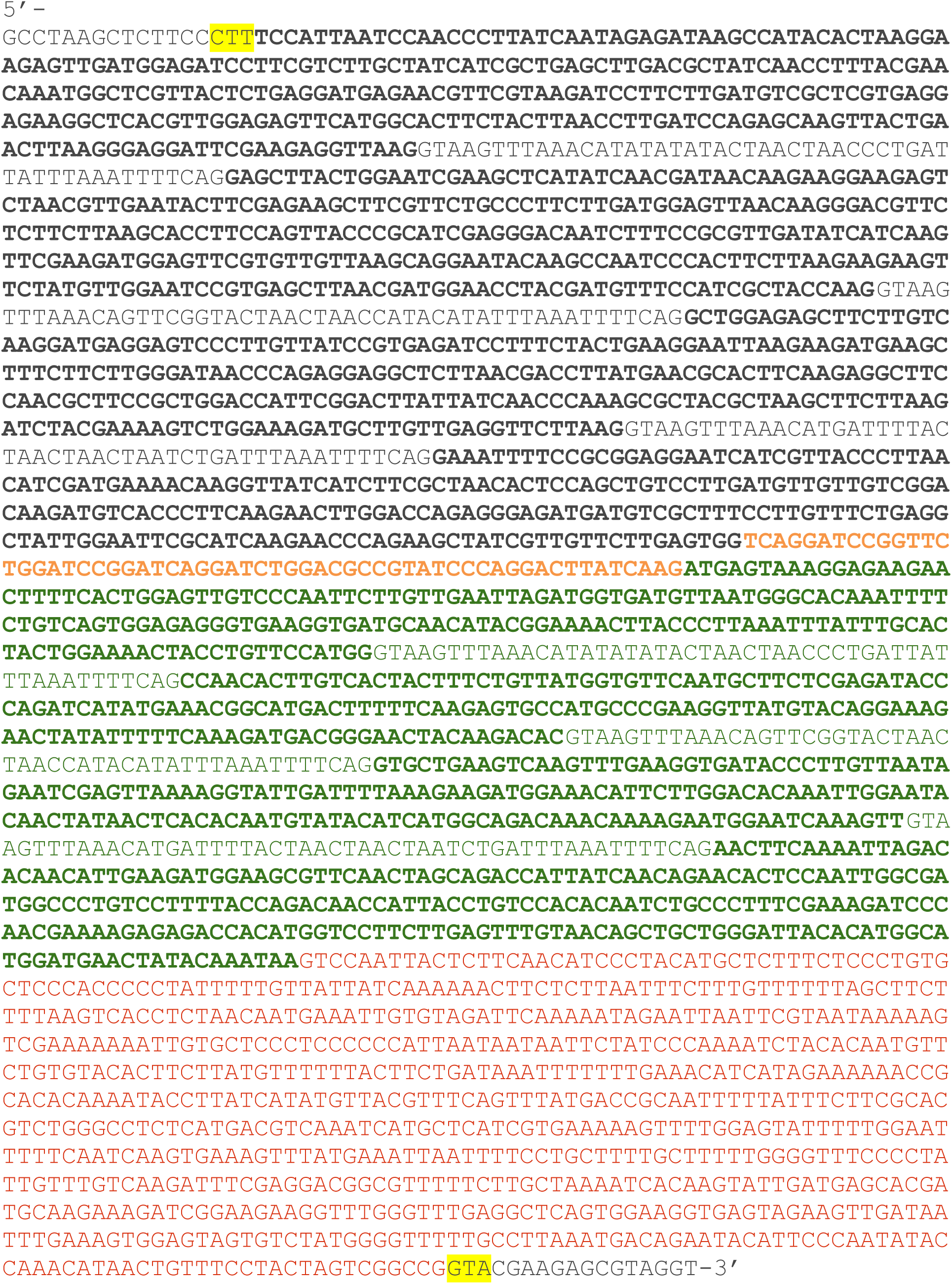
The synthesized gene block GEM sequence inserted into the plasmid. Overhangs produced by SapI are highlighted in yellow. The codon-optimized reading frame is bolded, with the *Pyrococcus furiosus* encapsulin 2E0Z protein encoded sequence in black, a 60bp linker in orange, and eGFP in green and introns are not bolded. The unc-54 3’-UTR is in red.

**Table. S1.** Strain list.

| Strain | Genotype | Reference |
| --- | --- | --- |
| NM5179 | *JsTi1493[LoxP::mex-5p::FLP:SL2::mNeonGreen::rpl-28p::FRT::GFP::his-58::FRT3] IV* | (26) |
| UD803 | *ycSi4[pSL848; p<sub>ges-1</sub>::40nmGEM::eGFP::unc-54 3'UTR] IV* | This study |
| UD838 | *ycSi5[pSL850; p<sub>semo-1</sub>::40nmGEM::eGFP::unc-54 3'UTR] IV* | This study |
| CB3440 | *anc-1(e1873) I* | (46) |
| UD573 | *anc-1(yc52[ΔKASH]) I* | (19) |
| UD590 | *anc-1(yc61[Δ6RPs]) I* | (19) |
| UD620 | *anc-1(yc71[ΔTK]) I* | (19) |
| VC40007 | *anc-1(gk109018[W621\*]) I* | (47) |
| UD848 | *ycSi4 IV; anc-1(e1873) I;* | This study |
| UD978 | *ycSi5 IV; anc-1(e1873) I* | This study |
| UD1036 | *ycSi4IV; anc-1(yc52[ΔKASH]) I* | This study |
| UD1035 | *ycSi5 IV; anc-1(yc52[ΔKASH]) I* | This study |
| UD1029 | *ycSi4 IV; anc-1 (yc61[Δ6RPs]) I* | This study |
| UD1032 | *ycSi5 IV; anc-1 (yc61[Δ6RPs]) I* | This study |
| UD1025 | *ycSi4 IV; anc-1(yc71[ΔTK]) I* | This study |
| UD1027 | *ycSi5 IV; anc-1(yc71[ΔTK]) I* | This study |
| UD1030 | *ycSi4 IV; anc-1(gk109018[W621\*]) I* | This study |
| UD1037 | *ycSi5 IV; anc-1(gk109018[W621\*]) I* | This study |
| UD756 | *ycSi2(pSL845[p<sub>semo-1</sub>::mKate2::tram-1]) IV* | This study |
| UD1039 | *ycSi2 IV; anc-1(e1873) I* | This study |
| UD1053 | *ycEx308[pSL915(p<sub>sur-5</sub>::gfp)]* | This study |
| UD1058 | *ycEx308; anc-1(e1873) I* | This study |
| UD863 | *yc113 IV; ycSi2 IV; JuEx5375* | This study |
| UD1063 | *yc113 IV; ycSi2 IV; JuEx5375; anc-1(e1873) I* | This study |
| UD522 | *ycEx249[p<sub>col-19</sub>::gfp::lacZ, P<sub>myo-2</sub>::mCherry]* | (48) |

**Table. S2.**
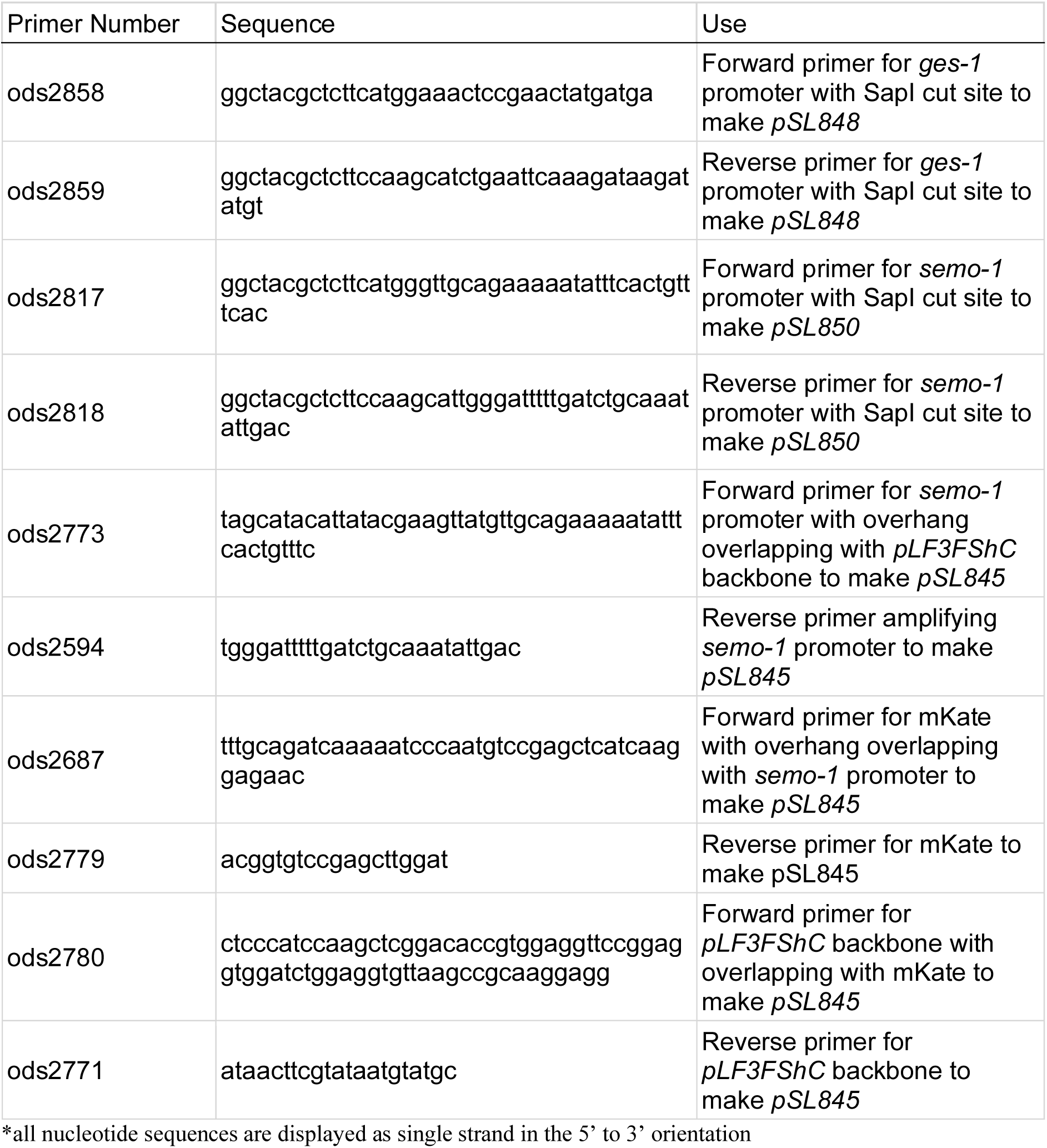
Primer list.

**Movie. S1.** Representative clip of a *C. elegans* young adult carrying the transgene *ycSi4[pSL848; pges-1::40nmGEM::eGFP::unc-54 3’UTR] IV*, showing GEMs movement in the intestine. GEMs are pseudocolored in green to highlight their dynamics.

**Movie. S2.** Representative clip of GEMs movement in the intestine of an *anc-1(e1873)* mutant *C. elegans* young adult carrying the transgene *ycSi4[pSL848; pges-1::40nmGEM::eGFP::unc-54 3’UTR] IV*. GEMs are pseudocolored in green to highlight their dynamics.

**Movie. S3.** Representative clip of GEMs movement in the hypodermis of a wild-type *C. elegans* young adult carrying the transgene *ycSi5[pSL850; psemo-1::40nmGEM::eGFP::unc-54 3’UTR] IV*. GEMs are pseudocolored in green to highlight their dynamics.

**Movie. S4.** Representative clip of GEMs movement in the hypodermis of an *anc-1(e1873)* mutant *C. elegans* young adult carrying the transgene *ycSi5[pSL850; psemo-1::40nmGEM::eGFP::unc-54 3’UTR] IV*. GEMs are pseudocolored in green to illustrate their dynamics.

**Movie. S5.** Representative movie of the hypodermal ER in a *C. elegans* young adult carrying the transgene *ycSi2(pSL845[Psemo-1::mKate2::tram-1]) IV* to mark the ER. The ER is pseudocolored in magenta.

